# Analyzing climate variations on multiple timescales can guide Zika virus response measures

**DOI:** 10.1101/059808

**Authors:** Á.G. Muñoz, M. C. Thomson, L. Goddard, S. Aldighieri

## Abstract

**Background:** The emergence of Zika virus (ZIKV) as a public health emergency in Latin America and the Caribbean (LAC) occurred during a period of severe drought and unusually high temperatures. Speculation in the literature exists that these climate conditions were associated with the 2015/2016 El Niño event and/or climate change but to date no quantitative assessment has been made. Analysis of related *flaviviruses* –such as dengue and chikungunya, which are transmitted by the same vectors– suggests that ZIKV dynamics is sensitive to climate seasonality and longer-term variability and trends. A better understanding the climate conditions conducive to the 2014-2016 epidemic may permit the development of climate-informed short- and long-term strategies for ZIKV prevention and control.

**Results:** Using a novel timescale-decomposition methodology, we demonstrate that extreme climate anomalies observed in most parts of South America during the current epidemic are not caused exclusively by El Niño or climate change –as speculated–, but are the result of a particular combination of climate signals acting at multiple timescales. In Brazil, the heart of the epidemic, we find that dry conditions present during 2013-2015 are explained primarily by year-to-year variability superimposed on decadal variability, but with little contribution of long-term trends. In contrast, the extreme warm temperatures of 2014-2015 resulted from the compound effect of climate change, decadal and year-to-year climate variability.

**Conclusions:** ZIKV response strategies adapted for a drought context in Brazil during El Niño 2015/2016 may need to be revised to accommodate the likely return of heavy rainfall associated with the probable 2016/2017 La Niña. Temperatures are likely to remain warm given the importance of long term and decadal scale climate signals.

## Background

It has been postulated that the 2015/2016 El Niño-Southern Oscillation and long-term climate change have contributed to the recent emergence of Zika virus (ZIKV) in Latin America and the Caribbean (LAC) [1]. While plausible, analysis on the climate-ZIKV interaction is constrained by the recent arrival of the virus in LAC and hence lack of historical time series of epidemiological data [2] and the diverse nature of prior epidemics across the globe [3]. Evidence to date suggests ZIKV is principally transmitted globally and in LAC by the container breeder mosquito *Aedes aegypti* [4]. Because of its recent rapid spread, *Ae. albopictus,* alongside other *Aedes spp.,* has been identified as a minor vector but one with significant transmission potential for the future [5]. Although ZIKV transmission depends on several factors including human behaviour, it is well established that the associated vectors are sensitive to variations in environmental temperature and rainfall, and weather-based early warning systems for the related dengue virus have been suggested in different regions of the world [6,7,8]. Temperature is a significant driver of the development rates of juvenile mosquito vectors and adult feeding/egg laying cycles, along with the length of extrinsic incubation period and viral replication of arboviruses [8,9,10,11]. Both excess rainfall and drought have been implicated in the creation of breeding sites for *Aedes* vectors of ZIKV, and associated epidemics of dengue and chikungunya. Heavy rainfall may result in the development of outdoor breeding sites in a wide range of artificial containers [12], whereas droughts may also encourage changes in human behaviour to water storage that results in increases in domestic breeding sites for *Aedes spp.* [13]

The climate at any location varies from its historical average on a number of time scales, including natural year-to-year and decadal (10-20 year) variations as well as long-term trends, the latter by construction compatible with anthropogenic climate change signals [14]. The magnitude or persistence of the climate variations may enhance or decrease epidemic potential in the region. In order to better understand how much of the total variance in rainfall and temperature is explained by different timescales, and how those variations connect to recent conditions that are connected in space and time to the emergence of ZIKV in LAC, we analyze how anomalies over time can be approximately attributed to variations in climate drivers at different timescales – an analysis referred to as “timescale decomposition” [14, 15]. This methodology filters the associated anomalies of a climate time series into three components: the inter-annual (year-to-year), decadal and long-term trend signals. The analysis shows how important each timescale is for explaining the entire historical climate signal observed in any particular location.

As indicated, the absence of long time series of ZIKV transmission indices or cases prohibits a formal statistical assessment of the climate-ZIKV linkage including the epidemiological impact of the 2015’s climate on the epidemic. However, our study is based on the premise that climate is likely an important driver of seasonal, inter-annual and longer-term trends in ZIKV transmission given that 1) temperature impacts the development rates of related arbovirus’ and the known vectors, and 2) droughts or excess of rainfall influence vector breeding sites either directly or through changes in human behavior. We therefore focus our analysis on the particular contributions of climate signals at multiple timescales to rainfall and temperature in order to support the development of climate-informed short- and long-term strategies for ZIKV prevention and control [14].

## Data Description

We chose two sources of climate data for our analysis as no single data set included the entire time of interest. Timescale decomposition (Figures 1 and 2) was undertaken using the most up-97 to-date long-term (1901-2014) rainfall and temperature data from University of East Anglia’s Climate Research Unit, product version 3.23 (CRUv3.23, 0.5-deg resolution) [16]. Recent annual temperature and rainfall anomalies (2013-2015, Figure 3) were computed using the Climate Prediction Center’s Monthly Global Surface Air Temperature Data Set (0.5 deg) [17] and Rainfall Unified Data Set (0.5 deg) [18], respectively. Years 1979-2000 were used to compute the normal for Figure 3.

Time series, maps and data are freely available in the IRI’s Timescale Decomposition Maproom [19] and the Latin American Observatory’s Climate and Health Maproom [20,21], and can be obtained for any region in the world having long enough quality-controlled records. For details, see [15].

## Analyses

The 20th Century decomposition for annual rainfall totals (Figure 1ABC) and annual mean temperature (Figure 1DEF) signals in LAC shows sharp differences in the variability explained by each timescale. The part of South America delimited by the black box in Figure 1 exhibits the highest number of reports associated with typical arbovirus vectors [22] and Zika cases [3], and thus it was selected for further analysis. On average, our results for the selected region indicate that the portion of variance in rainfall associated with the climate change signal is basically nil (Figure 1A), whereas that for the inter-annual component is about 60%-90% throughout the region (Figure 1C). Also, the decomposition reveals that all three timescale components for surface air temperature are important (Figure 1DEF).

The temperature long-term trend signal is particularly important along the southeastern regions of Brazil (Figure 1D). The decadal signal is in general more important for temperature than for rainfall in the region, the contribution to precipitation being higher along the coast (20%-30%, Figure 1B), whereas for surface air temperature the highest decadal component is found in the Amazon (~50%, Figure 1E). Inter-annual variations for surface temperature show values over 30% in most locations, with a local maximum in Northeastern Brazil that explains at least 60% of the variability (Figure 1F).

Results are similar for the region of interest when particular seasons are considered [19, 21]: for rainfall the most important scales are inter-annual and decadal, while for air surface temperature the three timescales share similar importance, while locally one timescale may exhibit greater importance than the others.

**Figure 1.**
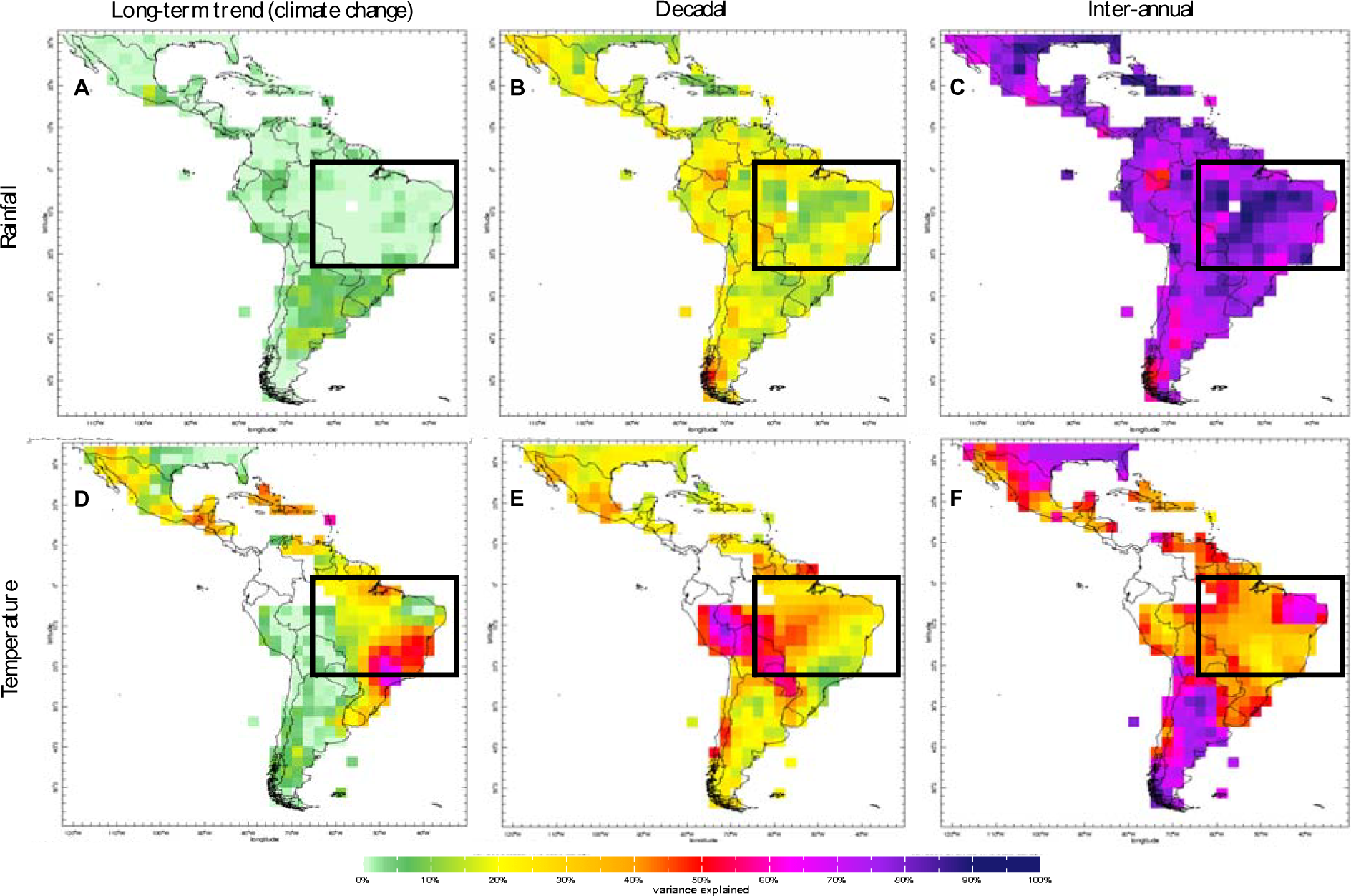
Timescale decomposition for annual precipitation (A,B,C) and air temperature (D,E,F), sketching the total explained variance for the long-term trend (A,D), decadal (B,E) and inter-annual variability (C,F) signals. Grid points in white indicate places where the lack of data would degrade the analysis and thus the corresponding signal has been removed by the screening process [15]. Analysis focuses in the region delimited by the black box (see main text).

We performed a complementary analysis for the average climate over the boxed region of interest (Figure 2). When summed, the specific contributions explain the observed anomalies for each particular year. Our results show that a positive superposition between the rainfall inter-143 annual and decadal signals and all three temperature components (climate change, decadal and inter-annual) is key to understand the recent behaviour of the climate in the region. This collection of drivers was responsible for the particularly warmer- and drier-than-normal conditions present in the region during the last few years. The unprecedented positive temperature anomalies that started in the 1990s are consistent with the positive sign of the decadal component for that period, combined with the contributions of the long-term trend and inter-annual variability.

**Figure 2.**
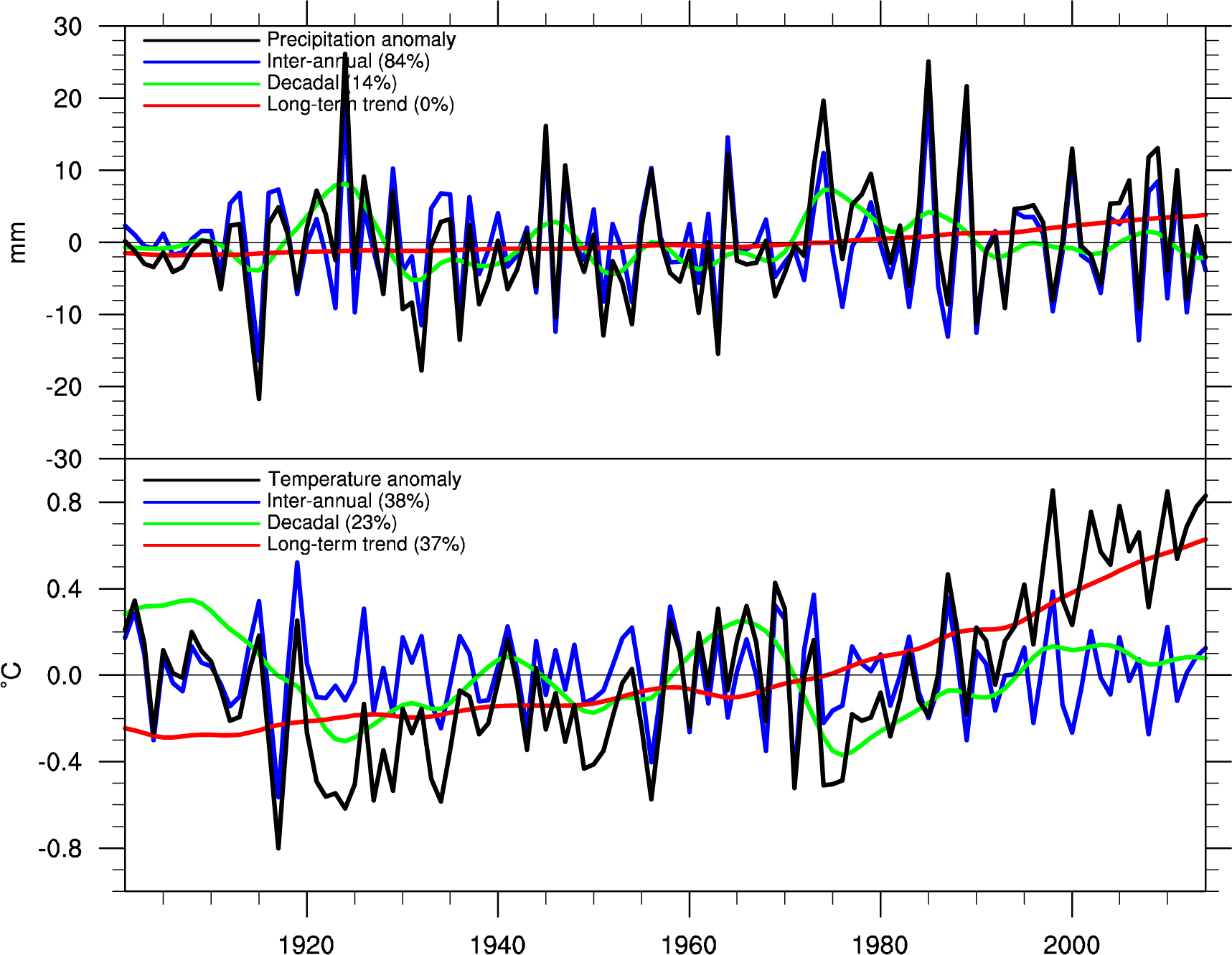
Timescale decomposition for annual anomalies in the 1901-2014 period (black curves; rainfall in the top panel, temperature in the bottom panel) averaged over the region indicated in Figure 1 (black box). The anomalies correspond to the superposition of the long-term trend (in red), the decadal signal (in green) and the inter-annual variability signal (in blue). Contribution of each timescale to the total explained variance is shown in parenthesis.

The spatial patterns for both temperature and rainfall anomalies in LAC were fairly similar in 2014 and 2015 (Figure 3), which were, at their respective terminus, the hottest years on record [23,24]; 2015 marked also the start of one of the three most intense El Niño events on record. In terms of temperature anomalies, the year 2013 was normal for most part of LAC, although the warming pattern in the Amazon that extended through the study region in the following years was already present. A similar claim can be made for the 2013 annual rainfall anomalies in the Amazon: the general drier-than-normal signal exhibited in 2014 and 2015 was already evolving then. Similar anomaly patterns were present in other countries too; for example, warmer-and-164 drier-than-normal conditions were observed in regions of Colombia, Venezuela, Ecuador and Puerto Rico, that also have been impacted by the ZIKV epidemic.

**Figure 3.**
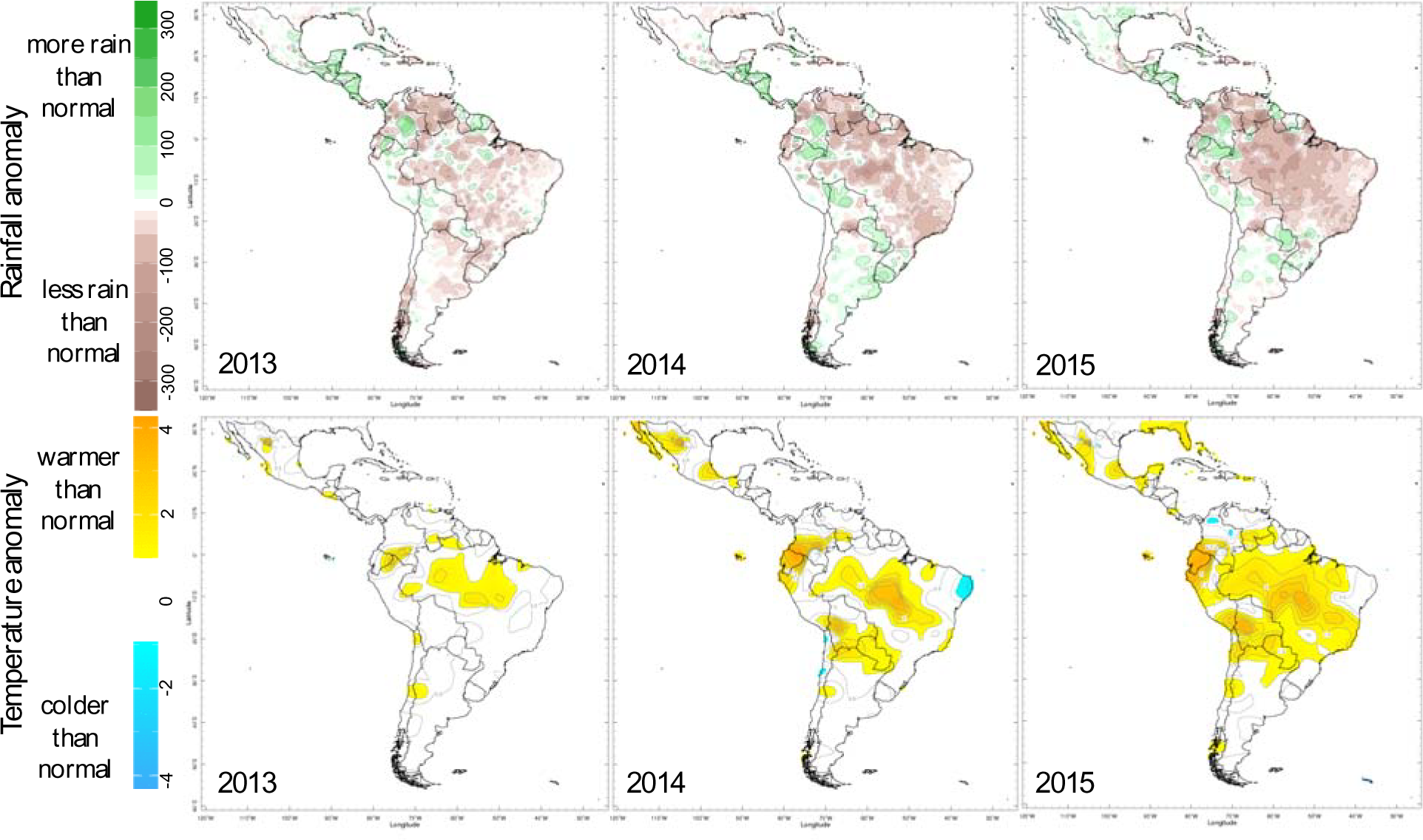
Annual temperature (upper row, in ^o^C) and rainfall (lower row, in mm) anomalies for 2013-2015. White indicates near-normal values.

## Discussion

The warming observed in 2014-15 is therefore an outcome of positive temperature anomalies at the year-to-year and decadal timescale superimposed on a long-term warming trend. This superposition of timescales may have helped to set the climate scenario for local ZIKV transmission via *Ae. aegypti* and other, less significant, vectors [4]. These patterns have also been observed during the first few months of 2016, although some rainfall anomalies have been changing as the year has progressed.

As of August 2016, seasonal forecasts of sea surface temperatures suggest about 55% probability of a La Niña event to be in place later this year [25], which is significantly higher than the corresponding climatological threshold (~35% for the same period). La Niña events typically lead to wetter than average conditions over the northern part of Brazil and Northern South America [26]. Since precipitation in this region is dominated by inter-annual variability, climate drivers at longer timescales are not likely to offset that response to La Niña. For temperature, the tropics tend to be relatively cooler during La Niña events, particularly relative to El Niño. However, given the comparable magnitude of decadal variability, which currently appears to be in a warm phase, and the strength of the long-term trend, warmer than average temperatures are still the most likely outcome over the coming year.

The characterization of year-to-year variability and longer term climatic trends is important for strategic ZIKV outbreak preparedness activities in LAC and across the border into the USA. For countries where variability and short and long-term trends are in part predictable, climate information could support the planning of prevention and control activities for different high risk areas, such as training of personnel in different aspects of the outbreak early warning and response system [27].

For example, the response strategies for ZIKV vector control in a warm and dry year, where high levels of water storage provide domestic breeding sites, may need revision in a wet year when outdoor breeding sites may be more common. Current speculation about the climate drivers that may impact ZIKV transmission (see for example [1]) are based on plausible assumptions regarding the dynamics of the disease but lack an in-depth understanding of the climate. However, using climate knowledge to improve health outcomes must be based on an understanding of the climate system itself and its interaction at multiple spatial and temporal scales. The timescale decomposition approach [15] used here allows a robust assessment of complex climate components to be made for any time period, season and region [19,21]. It provides a basis for considering climate as a resource to decision-maker efforts, not only for ZIKV, but also for other vector-borne diseases like chikungunya and dengue.

## Methods

Timescale decomposition consists of screening the individual gridbox values for filled data and for very dry seasons and regions; detrending in order to extract slow, trend-like changes; and filtering, to separate high and low frequency components in the detrended data. Detrending involves regressing the local time series on multi-model global surface air temperature from the Twentieth Century Climate in Coupled Models [28] and low-pass filtering. The decadal component is obtained through low-pass filtering of the residual, using an order-five Butterworth filter with half-power at a period of 10 years, while the inter-annual component is computed as the difference between the residual from the detrending step and the decadal signal [15]. For additional details, see the International Research Institute for Climate and Society (IRI) Timescale Decomposition Maproom [19].

For the maps in Figure 1, data are processed gridbox by gridbox, meaning that results in adjacent gridboxes are not compared or combined. For the graph of the regional timeseries (Figure 2), averaging over gridboxes is performed prior to the decomposition. Total explained variance for each component is computed for the area-averaged time series, and not as averages of the spatial variance maps.

## Availability and requirements

- Project name: Climate and Health Maproom
- Project home page: http://iridl.ldeo.columbia.edu/maproom/Health/index.html and http://datoteca.ole2.org/maproom/Sala_de_Salud-Clima/index.html.es
- Archived version: http://doi.org/10.1029/2011EO450001
- Operating system(s): Platform independent
- Programming language: Ingrid
- Other requirements: none
- License: Open Database License (ODbL) v1.0

## Abbreviations

LAC: Latin America and the Caribbean
ZIKV: Zika virus

## Availability of supporting data

Data and figures supporting the results of this research are freely available online in the IRI’s Timescale Decomposition Maproom [19] and the Latin American Observatory’s Climate and Health Maproom [20,21].

## Declarations

### Acknowledgements

We acknowledge the assistance of Rémi Cousin and Xandre Chourio with IRI’s and Latin American Observatory’s Timescale Decomposition Maproom datasets, as well as Xiaosong Yang’s and Catherine Vaughan’s comments on the manuscript. Muñoz was supported by NOAA/OAR under the auspices of the National Earth System Prediction Capability (National ESPC).

## Author’s Contributions

Á.G.M., M.C.T. and S.A established the concept of the study. Á.G.M. obtained the data. Á.G.M., M.C.T and L.G undertook the analysis and interpretation of results. Á.G.M., M.C.T and L.G. drafted the manuscript. All authors critically reviewed and revised the manuscript and agreed the final submission.

